# Allosteric MAPKAPK2 Inhibitors Improve Plaque Stability in Advanced Atherosclerosis

**DOI:** 10.1101/2021.01.12.426264

**Authors:** Lale Ozcan, Canan Kasikara, Arif Yurdagul, George Kuriakose, Brian Hubbard, Michael H. Serrano-Wu, Ira Tabas

## Abstract

Atherosclerotic vascular disease resulting from unstable plaques is the leading cause of morbidity and mortality in subjects with type 2 diabetes (T2D), and thus a major therapeutic goal is to discover T2D drugs that can also promote atherosclerotic plaque stability. Genetic or pharmacologic inhibition of mitogen-activated protein kinase-activated protein kinase-2 (MK2) in obese mice improves glucose homeostasis and enhances insulin sensitivity. We developed two novel orally active small-molecule inhibitors of MK2, TBX-1 and TBX-2, and tested their effects on metabolism and atherosclerosis in high-fat Western diet (WD)-fed *Ldlr*^-/-^ mice. *Ldlr*^-/-^ mice were first fed the WD to allow atherosclerotic lesions to become established, and the mice were then treated with TBX-1 or TBX-2. Both compounds improved glucose metabolism and lowered plasma cholesterol and triglyceride, without an effect on body weight. Most importantly, the compounds decreased lesion area, lessened plaque necrosis, and increased fibrous cap thickness in the aortic root lesions of the mice. Thus, in a preclinical model of high-fat feeding and established atherosclerosis, MK2 inhibitors improved metabolism and also enhanced atherosclerotic plaque stability, suggesting potential for further clinical development to address the epidemic of T2D associated with atherosclerotic vascular disease.

## INTRODUCTION

The prevalence of type 2 diabetes (T2D) is becoming a significant public health problem owing to the worldwide obesity epidemic [1]. Patients with T2D have a two- to four-fold increased risk of developing atherosclerotic cardiovascular disease (CVD), which accounts for ~40% of all T2D mortality [2, 3]. Reducing the incidence and severity of CVD is a key unmet medical need in T2D patients. Over the past decade, many anti-diabetic agents have been tested for cardiovascular safety [4]. Despite good glycemic control, most of these studies failed to show a significant CVD benefit [5]. Therefore, identification of common signaling pathways and molecular targets in distinct cell types that contribute to both T2D and CVD pathogenesis may lead to the development of novel diabetes therapies with proven CVD benefits.

We have previously shown that dysregulated calcium signaling in hepatocytes in obesity activates a calcium/calmodulin-dependent protein kinase II (CaMKII)?MK2 pathway, which then promotes hyperglycemia and insulin resistance via increasing hepatic glucose production and disrupting insulin receptor signaling [6–8]. Genetic inhibition of hepatic CaMKII or MK2 in obese insulin-resistant mouse models markedly lowers blood glucose and insulin resistance [7]. Further, treatment with an allosteric MK2 inhibitor called compound (cmpd) 28 was shown to improve glucose homeostasis and insulin sensitivity in obese mice [9]. The metabolic benefit of cmpd 28 treatment in this study was mechanistically “on-target” and, importantly, was additive with the current leading T2D drug, metformin. Consistent with these results, MK2 deletion has recently been shown to lower hyperglycemia and improve insulin tolerance in the low-dose streptozotocin-induced diabetic mouse model [10]. Interestingly, a different CaMKII-MK2–driven downstream pathway in atherosclerotic lesional macrophages promotes advanced atherosclerosis progression by impairing clearance of apoptotic cells (efferocytosis) and suppressing resolution of inflammation [11, 12].

The mechanisms linking T2D to CVD include insulin resistance, hyperglycemia, dyslipoproteinemia, and inflammation [13]. Our pre-clinical studies have shown that inhibition of the CaMKII?MK2 pathway ameliorates all of these processes [7, 9], and germline deletion of MK2 in *Ldlr*^-/-^ mice was shown to suppress atherosclerosis, but effects on plaque stability and glucose metabolism was not reported in that study [14]. Using two new orally available MK2 inhibitors, TBX-1 and TBX-2, we now report that MK2 inhibitor treatment of *Ldlr*^-/-^ mice with established atherosclerosis improves metabolism and features of unstable plaques.

## MATERIALS AND METHODS

### Synthesis of MK2 inhibitors

TBX-1 and TBX-2 were synthesized as described in the patent literature [15]. All solvents and reagents were obtained from commercial sources and used without further purification. NMR spectra were obtained on a Bruker Neo 400M spectrometer operating at 400 MHz. Chemical shifts are reported in parts per million (δ) from the tetramethysilane resonance in the indicated solvent. LC-Mass spectra were taken with Agilent 1260-6125B single quadrupole mass spectrometer using a Welch Biomate column (C18, 2.7 um, 4.6*50 mm) eluting with a mixture of solvents A (ACN with 0.05% trifluoroacetic acid) and B (Water with 0.05% trifluoroacetic acid) using a gradient elution. Detection was by DAD (254 nm and 210 nm). Ionization was by ESI.

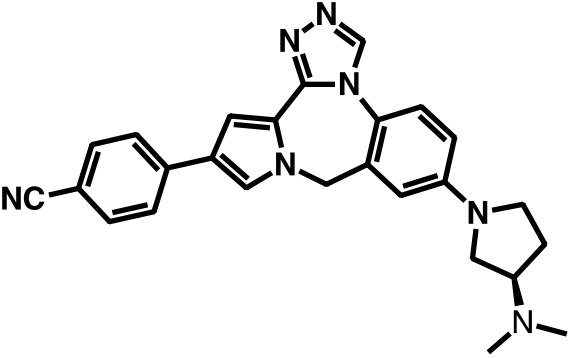
(R)-4-(7-(3-(Dimethylamino)pyrrolidin-1-yl)-9H-benzo[e]pyrrolo[1,2-a][1,2,4] triazolo[3,4-c][1,4]diazepin-12-yl)benzonitrile (TBX-1) ESI [M+H]^+^ = 436; ^1^H NMR (400 MHz, CDCl3) δ 8.53 (s, 1H), 7.62 – 7.53 (m, 4H), 7.27 (d, 1H, *J* =7.6 Hz), 7.24 (d, *J* = 1.9 Hz, 1H), 7.21 (d, *J* =1.9 Hz, 1H) 6.58-6.54 (m, 2H), 4.99 (s, 2H), 3.59 – 3.44 (m, 2H), 3.36 (td, *J* = 9.6, 6.9 Hz, 1H), 3.21 (t, *J* = 8.4 Hz, 1H), 2.98 – 2.83 (m, 1H), 2.34 (s, 6H), 2.30 – 2.22 (m, 1H), 2.08 – 1.91 (m, 1H).

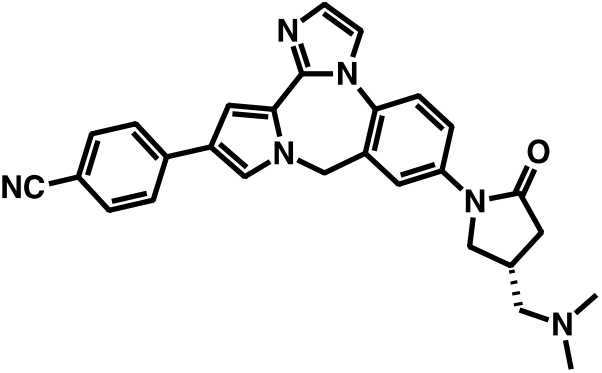
(R)-4-(7-(4-((Dimethylamino)methyl)-2-oxopyrrolidin-1-yl)-9H-benzo[e]imidazo[2,1-c]pyrrolo[1,2-a][1,4]diazepin-12-yl) benzonitrile (TBX-2) ESI [M+H]^+^ = 463; ^1^H NMR (400 MHz, MeOH-*d4*) δ 7.97 (d, *J* = 2.5 Hz, 1H), 7.78 (dd, *J* = 8.8, 2.5 Hz, 1H), 7.74 – 7.67 (m, 3H), 7.66 – 7.62 (m, 2H), 7.58 (d, *J* = 8.8 Hz, 1H), 7.53 (d, *J* = 1.9 Hz, 1H), 7.25 (d, *J* = 1.3 Hz, 1H), 7.05 (d, *J* = 1.9 Hz, 1H), 5.18 (s, 2H), 4.05 (dd, *J* = 9.7, 7.8 Hz, 1H), 3.76 – 3.67 (m, 1H), 2.83 – 2.71 (m, 2H), 2.51 (dd, *J* = 12.3, 7.7 Hz, 1H), 2.47 – 2.37 (m, 2H), 2.30 (s, 6H).

### Mouse Experiments

Eight-week-old male *Ldlr*^-/-^ mice on the C57BL/6J background were purchased from Jackson Laboratory (stock # 002207) and fed a Western-type Diet (WD) for 16 weeks (Envigo, TD.88137; 21.2% fat, 0.2% cholesterol, 48.5% carbohydrate, and 17.3% protein by weight). Between weeks 10-16 of feeding, the WD of the experimental group was supplemented with TBX-1 or TBX-2 (600 mg/kg diet) to achieve a dose of ~30 mg/kg body weight per day, whereas control mice remained on regular WD. The study included four separate cohorts, each comprised of 5 to 10 mice per group. The mice were randomly assigned to the different cohorts. Fasting blood glucose was measured in mice that were fasted for 5 h, with free access to water, using a glucose meter. Insulin tolerance test was performed in 5 h-fasted mice by assaying blood glucose at various times after injection of insulin (0.7 IU/kg body wt i.p.). Plasma insulin levels were measured in mice that were fasted for 5 h using an ultra-sensitive mouse insulin ELISA kit (Crystal Chem). The Columbia University IACUC provided ethical approval for these studies, and procedures were conducted in accordance with approved animal protocols.

### Pharmacokinetic studies

The pharmacokinetics of the compounds were evaluated in CD-1 mice by single oral and intravenous routes of drug administration. The mice were fasted overnight before oral-gavage dosing of the compounds. The solutions used for PK studies were prepared by dissolving the compounds in 5% Solutol HS 15 in water. Following oral or intravenous administration of compounds, blood samples were collected at 0.08, 0.25, 0.5, 1, 2, 4, 6, 8, 12, and 24 h after dosing. Blood samples were centrifuged to separate plasma, and compound concentrations were determined by LC-MS. Dosing was well tolerated following oral and i.v. administration of both compounds, and no abnormal behavior was observed.

### Plasma cholesterol and triglyceride measurements

Overnight fasting plasma was collected from mice via exsanguination from left-ventricular puncture. Total plasma cholesterol and triglyceride levels were measured using commercially available kits (Wako Diagnostics). Plasma lipoproteins were analyzed by running 200 μl of pooled plasma onto a fast protein liquid chromatography system consisting of 2 Superose 6 columns connected in series (Amersham Pharmacia Biotech) as described previously [16].

### Murine atherosclerotic lesion analysis

Total lesion area and necrotic area analysis were carried out on hematoxylin and eosin (H&E)-stained aortic root lesional cross sections as previously described [12]. Briefly, total lesion area (from internal elastic lamina to the lumen) and acellular/anuclear areas (negative for hematoxylin-positive nuclei) per cross section were quantified by taking the average of 6 sections spaced 30 μm apart beginning at the base of the aortic root. Necrotic areas were defined as anuclear areas based on absence of hematoxylin staining. Collagen staining was performed using picrosirius red as per the manufacturer’s instructions (Polysciences Inc.), and fibrous cap thickness was quantified from 3 distinct regions of the plaque and scored per lesion size as previously described [11]. For immunofluorescence staining, frozen sections were fixed in cold acetone for 10 minutes. After blocking for 1 hour in serum-free protein blocking buffer (DAKO, catalog X0909), sections were incubated overnight at 4°C with anti–phospho-hsp25 (1:200) and anti-hsp25 (1:200) antibodies, rinsed with PBS three times, incubated with fluorescently-labeled secondary antibodies for 2 hours, and counterstained with DAPI. Three slides, each with 2 sections, were assessed for each mouse.

### In situ efferocytosis assay

Experimentation was carried out as previously described [12]. In brief, acetone-fixed aortic root sections were incubated with TUNEL (Roche) followed by staining with anti-Mac2 (1:10,000, Cedarlane) to label macrophages. Apoptotic cells were then determined to be either macrophage-associated (colocalizing or juxtaposed with macrophages) or free (not associated with macrophages). Data were plotted as a ratio of associated-to free cells, which is a measure of efferocytosis [17].

### Primary hepatocytes

Primary mouse hepatocytes were isolated from 8- to 12-week-old mice as described previously [9] and cultured in media containing Dulbecco’s Modified Eagle’s Medium (DMEM) (Corning, #10-013-CV), 10% (vol/vol) heat-inactivated FBS, and 1X penicillin-streptomycin. After overnight incubation, the cells were washed twice with PBS, treated with the indicated concentrations of TBX-1 for 1 h, followed by treatment with TBX-1 and forskolin for 4 h in FBS-free media.

### Reagents and antibodies

Forskolin was from Sigma. Anti–phospho-hsp25 (CST, 2401) and anti-hsp25 (CST, 2442) antibodies were from Cell Signaling, and anti–β-actin antibody was from Abcam.

### Immunoblotting and quantitative RT-PCR

Immunoblot and quantitative RT-PCR (RT-qPCR) assays were conducted as previously described [9]. Total RNA was extracted from primary hepatocytes using the RNeasy kit (Qiagen). cDNA was synthesized from 2 mg total RNA using oligo (dT) and Superscript II (Invitrogen).

### Statistics

All results are presented as means ± SEM. P values were calculated using the Student’s t test for normally distributed data and Mann-Whitney rank sum test for non-normally distributed data.

## RESULTS

Our previous MK2 inhibitor study in obese mice used cmpd 28 [9]. However, the low oral bioavailability of cmpd 28 (BAV < 3%) precluded use of this tool compound in longer term animal experiments. Following a series of scaffold re-designs (to be described elsewhere), two MK2 inhibitors with greatly improved oral bioavailability were identified: TBX-1 (hMK2 IC_50_ = 67 nM) and TBX-2 (hMK2 IC_50_ = 19 nM). Similar to cmpd 28 [9], TBX-1 inhibited forskolin-induced *G6pc* mRNA in a dose-dependent manner (**Fig S1A**). The improved pharmacokinetic profiles of TBX-1 and TBX-2 were affirmed by intravenous (i.v.) and oral (p.o.) administration routes in normal chow-fed mice (**Table 1**). The plasma half-lives of TBX-1 and TBX-2 were 2.7 and 3.1 hours, respectively. Both compounds displayed clearance and absorption profiles that were suitable for oral delivery, making them attractive small molecule inhibitors to study the role of MK2 in advanced atherosclerotic lesion formation in mice.

**Table 1.**
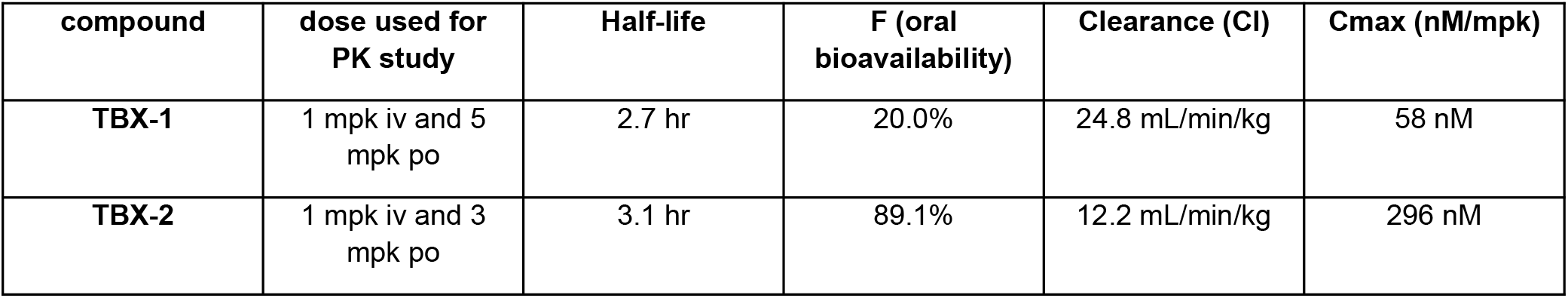
In vivo pharmacokinetic evaluation of compounds TBX-1 and TBX-2. Mice received indicated doses of compounds via oral or intravenous route. Blood samples were collected at 0.08, 0.25, 0.5, 1, 2, 4, 6, 8, 12, 24 h after dosing, and compound concentrations were determined by LC-MS.

To determine the therapeutic potential of MK2 inhibitors and provide clinically relevant findings, we tested whether they could improve features of plaque stability when administered after plaques were established. For this purpose, *Ldlr*^-/-^ mice were first fed a Western-type diet (WD) for 10 weeks to induce atherosclerotic plaque development. Based on the pharmacokinetic profile of the compounds, we supplemented the WD with 600 mg/kg TBX-1 or TBX-2 to achieve a dose of ∼30 mg/kg body weight per day. Half of the mice were continued for 6 additional weeks on the MK2 inhibitor supplemented diets, while the other half continued to receive WD alone (control). Neither treatment was associated with overt signs of toxicity, e.g., body weights, food intake, and activity levels were similar to those of control mice. As predicted, inhibitor treatment efficiently reduced the phosphorylation of the MK2 substrate, heat-shock protein (hsp) 25 in liver, as demonstrated with TBX-1 (**Fig S1B**), indicating successful targeting of MK2 activity in liver. As with cmpd 28, both inhibitors improved glucose homeostasis and enhanced insulin sensitivity (**Fig 1**). Both compounds also lowered plasma levels of total cholesterol (TC) and triglyceride (**Fig 2A, B, D, and E**). Fast protein liquid chromatography analysis of the plasma showed that very-low-density lipoprotein (VLDL)-cholesterol and low-density lipoprotein (LDL)-cholesterol were reduced after treatment with TBX-1 and TBX-2 (**Fig 2C** and **Fig 2F**). Analysis of white blood cells in the blood revealed no differences in the number of circulating leukocytes or monocytes (**Fig S2**).

**Figure 1.**
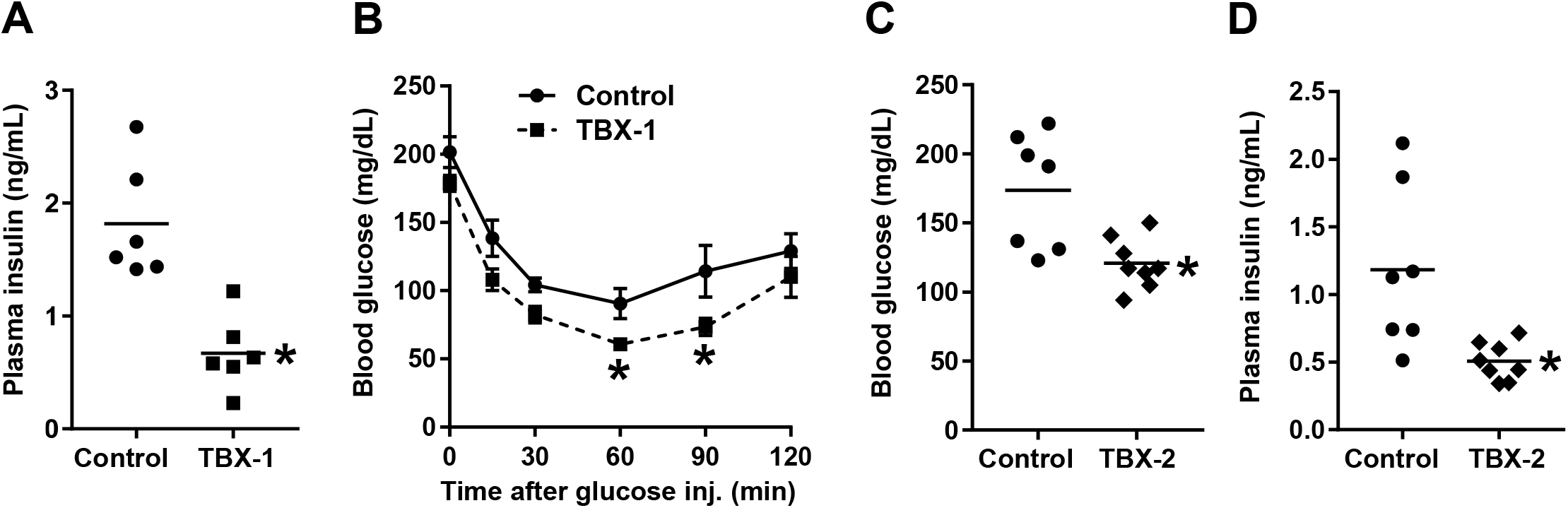
MK2 inhibitor treatment improves glucose homeostasis in WD-fed *Ldlr*^-/-^ mice. 8-week-old *Ldlr*^-/-^ mice were placed on WD for 10 weeks. The mice were continued for 6 additional weeks on WD alone (control) or on WD containing TBX-1 or TBX-2. (**A-B**) 5-hour-fasting plasma insulin levels were measured, and an insulin tolerance test was conducted in a randomly selected subset of *Ldlr*^-/-^ mice treated withTBX-1 (n = 6 mice/group; mean ± SEM, p < 0.05). (**C-D**) 5-hour fasting blood glucose and plasma insulin levels were measured in a randomly selected subset of *Ldlr*^-/-^ mice treated with TBX-2 (n = 7-8 mice/group; mean ± SEM, p < 0.05).

**Figure 2.**
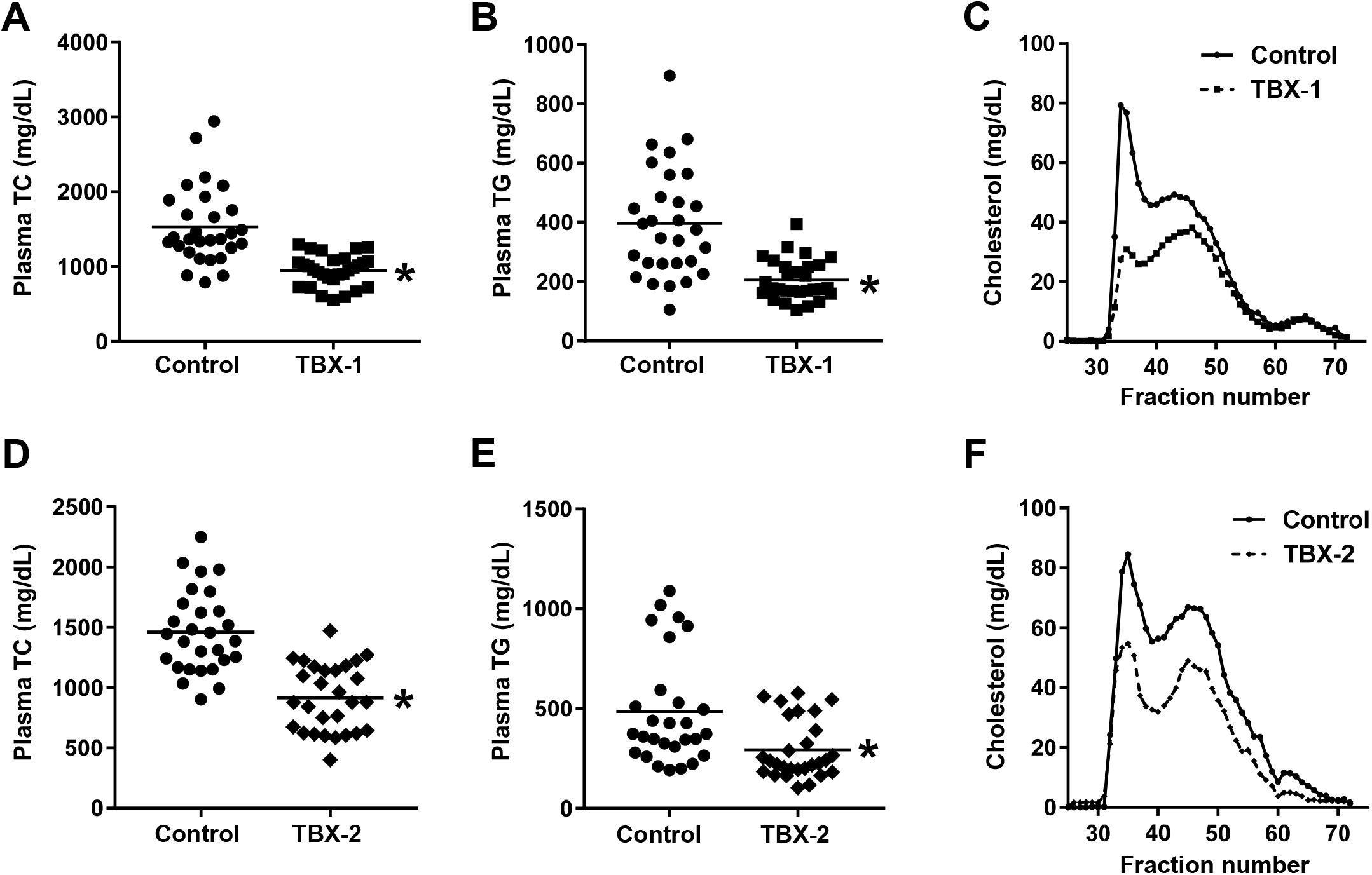
MK2 inhibitor treatment lowers plasma lipids in WD-fed *Ldlr*^-/-^ mice. Plasma total cholesterol (TC) and triglyceride (TG) levels were measured, and FPLC-separated lipoproteins were assayed for cholesterol, in *Ldlr*^-/-^ mice treated with TBX-1 (**A-C**) or TBX-2 (**D-F**) as in Figure 1 (n = 29 mice/group for TBX-1 and n = 28 mice/group for TBX-2; mean ± SEM, p < 0.05).

Turning to atherosclerotic lesion analysis the aortic root, we first documented that MK2 activity was reduced in the atherosclerotic plaques of inhibitor-treated mice, as phospho-hsp 25 levels were significantly reduced in the lesions of *Ldlr*^-/-^ mice treated with TBX-1 (**Fig 3A**). We then quantified the overall lesion area and necrotic area of the aortic root plaques and found that both treatments lowered total lesion area compared with untreated control mice (**Fig 3B** and **Fig 3D**). Moreover, total necrotic area was significantly reduced in mice treated with TBX-1 and TBX-2 (**Fig 3C** and **Fig 3E**).

**Figure 3.**
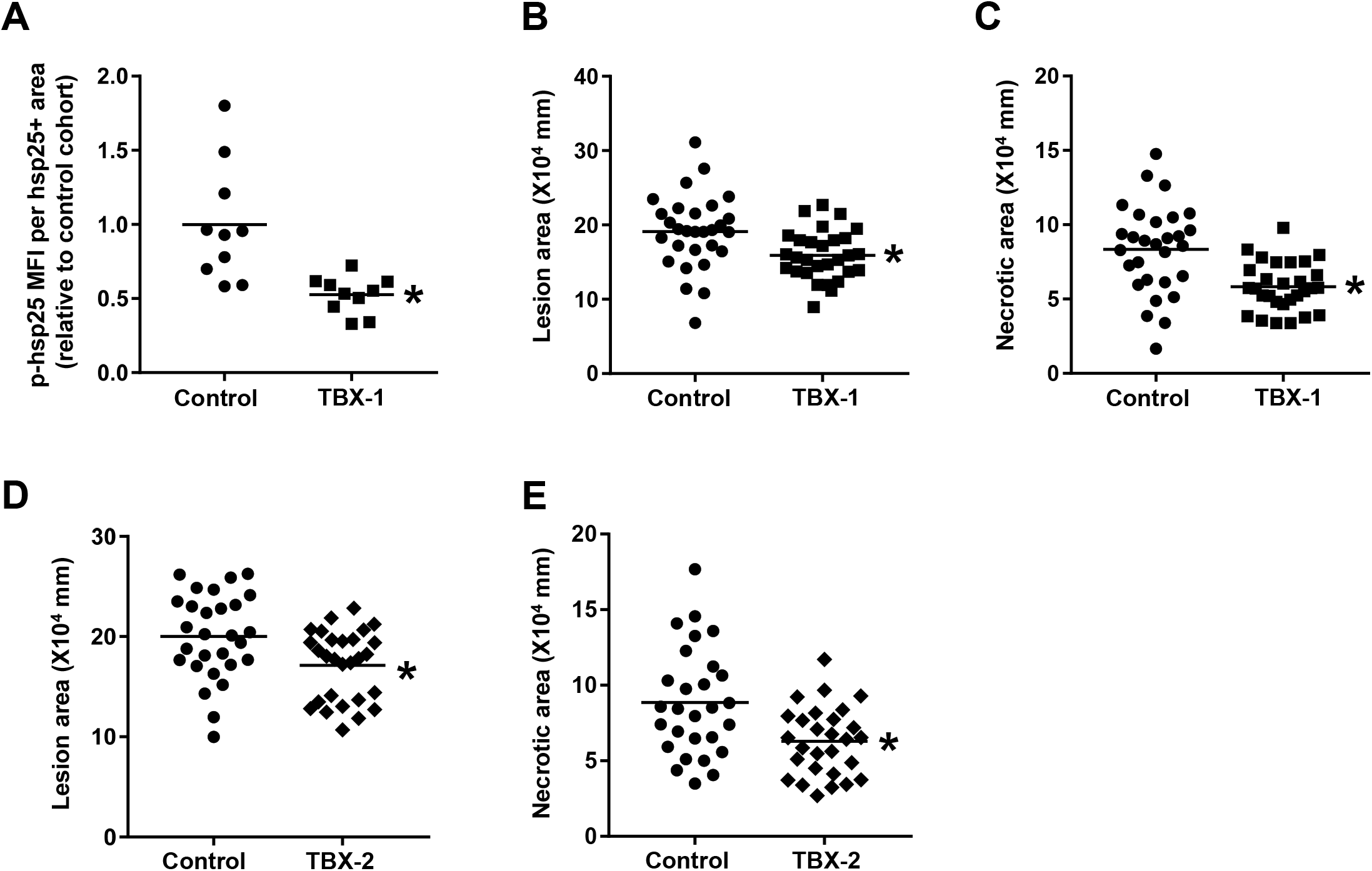
MK2 inhibitor treatment lowers p-hsp25, lesion size, and plaque necrosis in aortic root lesions of WD-fed *Ldlr*^-/-^ mice. Mice were treated as in Figure 1. (**A**) Aortic root sections from a randomly selected subset of *Ldlr*^-/-^ mice treated with TBX-1 were co-immunostained for p-hsp25 and total hsp25. Phospho-hsp25 staining was quantified as MFI within hsp25^+^ cells. Data are presented relative to the average value obtained from the control group (n = 10 mice/group; mean ± SEM, p < 0.05). (**B-E**) Total lesion area and necrotic area were quantified from the aortic root sections of *Ldlr*^-/-^ mice treated with TBX-1 (**B-C**) or TBX-2 (**D-E**) (n = 29 mice/group for TBX-1 and n = 28 mice/group for TBX-2; mean ± SEM, p < 0.05).

Impaired efferocytosis is a key driver of necrotic core formation in advanced atherosclerosis [18]. To determine whether MK2 inhibition improved lesional efferocytosis, we used an *in-situ* measure of efferocytosis in the lesions in which the ratio of macrophage-associated to free apoptotic cells is quantified. We found that both compounds significantly enhanced lesional efferocytosis, which could explain the decrease in plaque necrosis (**Fig 4A** and **Fig 4C**). Fibrous-cap thickness, which is another factor that promotes plaque stability [19], was significantly increased in TBX-1– and TBX-2–treated mice (**Fig 4B** and **Fig 4D**). Collectively, these data indicate that MK2 inhibition induces a more stable plaque phenotype with less plaque necrosis and thicker fibrous caps.

**Figure 4.**
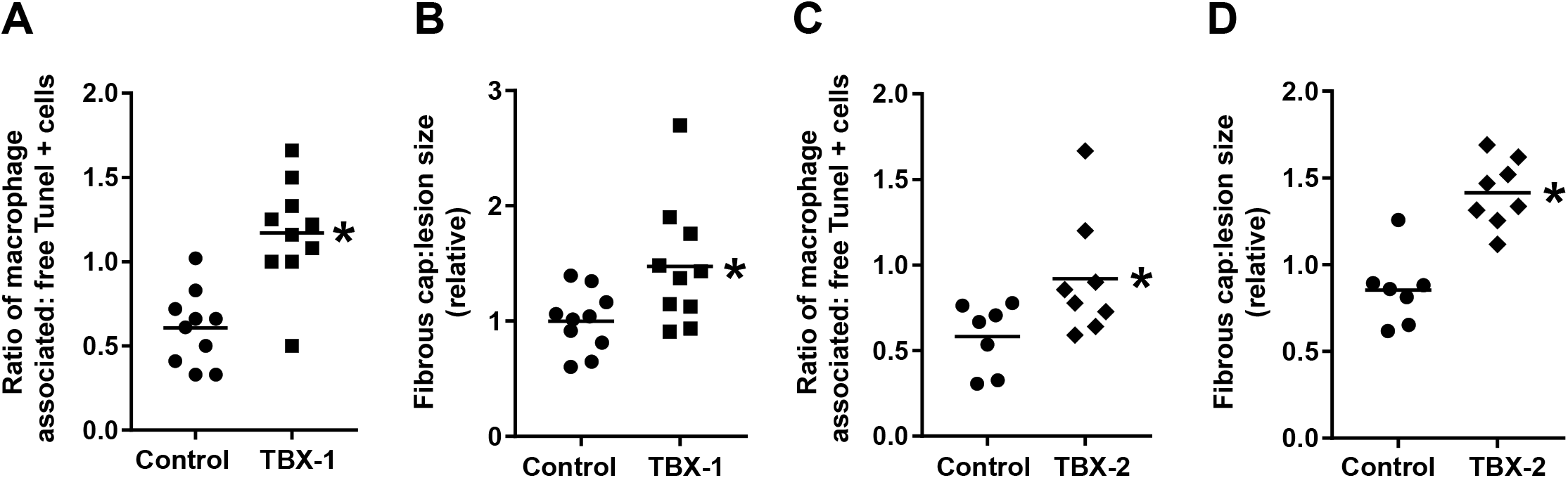
MK2 inhibitor treatment enhances fibrous cap formation and lesion efferocytosis in WD-fed *Ldlr*^-/-^ mice. Mice were treated as in Figure 1. (**A**) Lesional efferocytosis was quantified from aortic sections as the ratio of TUNEL^+^ cells associated with Mac2+ macrophages to free TUNEL^+^ cells in a randomly selected subset of TBX-1–treated *Ldlr*^-/-^ mice (n = 10 mice/group; mean ± SEM, p < 0.05). (**B**) Aortic root sections from a randomly selected subset of TBX-1–treated *Ldlr*^-/-^ mice were stained with picrosirius red, and fibrous cap was measured at the midpoint and shoulder regions of each lesion and quantified as the ratio of cap thickness to lesion size (n = 10 mice/group; mean ± SEM, p < 0.05). (**C-D**) As in (A-B) except that aortic sections from a randomly selected subset of TBX-2–treated mice were analyzed (n = 7-8 mice/group; mean ± SEM, p < 0.05).

## DISCUSSION

In response to diverse stimuli, MK2 is activated via p38 MAPK-mediated phosphorylation, and activated MK2 plays a key role in a variety of inflammatory and metabolic diseases, including heart disease and T2D [20]. Inhibition of MK2 activity causes a partial decrease in p38 via p38 destabilization and offers a safe therapeutic advantage over p38 inhibitors [21]. We recently reported that treatment of obese insulin-resistant mice with a small-molecule inhibitor of MK2 improves metabolism and lowers blood glucose [9]. As insulin resistance and hyperglycemia are linked to the development of clinically significant unstable atherosclerotic plaques [22], and activated MK2 in lesional macrophages can impair efferocytosis and resolution [11, 12], we hypothesized that MK2 could represent a promising therapeutic target to prevent unstable plaque formation in the setting of insulin resistance. In this study, we have identified two new orally active small-molecule MK2 inhibitors, TBX-1 and TBX-2, and tested their effect on advanced atherosclerosis progression of established plaques in high fat-fed *Ldlr*^-/-^ mice. We found that both compounds promoted features of plaque stability as evidenced by lower lesion area and plaque necrosis and increased fibrous cap thickness. The consistent biological effects elicited by two chemically distinct MK2 inhibitors provides strong support for the therapeutic benefit of MK2 inhibition to ameliorate chronic inflammation in CVD.

The ability of the compounds to lower circulating levels of atherogenic apolipoprotein B-containing lipoproteins undoubtedly contributes to their atheroprotective effect. Our previous work revealed that cmpd 28 treatment of DIO mice also resulted in a trend towards lowering of plasma lipids [9]. However, it is possible that other mechanisms could have contributed to the overall lesional phenotype. Chronic unresolved inflammation is a key feature of advanced atherosclerosis. In this regard, MK2 has been shown to stimulate many proinflammatory cytokines, and MK2-deficient mice are protected from LPS-induced endotoxic shock and inflammation-driven tumorigenesis [23, 24]. Further, MK2 directly phosphorylates and regulates the nuclear localization of 5-lipoxygenase (5-LOX), which limits the synthesis of proresolving molecules and favors proinflammatory mediator formation [25, 26]. Proresolving mediators enhance lesional efferocytosis and increase fibrous cap thickness in athero-prone mice [27], and so it is possible that a direct effect of MK2 inhibition on macrophage inflammatory/proresolving phenotype could have contributed to the improved plaque stability in the mice in the current study.

The possibility of a direct anti-atherosclerosis effect of the MK2 inhibitors, i.e., independent of lipid-lowering, is supported by a study in which genetic targeting of MK2 led to decreased lesion area in fat-fed *Ldlr*^-/-^ mice, as, curiously, plasma lipids were actually higher, not lower, in the *Mapkapk2*^-/-^ *Ldlr*^-/-^ mice used in this study [14]. The authors showed that MK2 activity is enriched in endothelial cells and macrophages in atherosclerotic lesions, and they provided evidence that the atheroprotective effects of MK2 targeting might be due to suppression of macrophage foam cell formation and decreased endothelium-mediated recruitment of monocytes into lesions. In that study, parameters of glucose metabolism and plaque stability were not reported. Although the difference in lipoprotein effects between that study and ours could be due to an MK2-independent mechanism of lipoprotein lowering by the class of compounds used here, another explanation could be the difference in study design. In the previous study, MK2 was completely absent before the high fat diet was begun. Moreover, germline gene targeting can sometimes lead to unexpected compensatory changes in the expression of other genes, leading to phenotypic changes not directly related to the targeted gene [28]. In our study, MK2 was inhibited, not completely deleted, 10 weeks after the high-fat diet was begun in mice with genetically intact MK2. Furthermore, given the high sequence identity (75%) between MK2 and MK3, it is reasonable to expect that both TBX-1 and TBX-2 inhibit MK2 and MK3 with similar efficiency, as the two molecules share a common pharmacophore [29, 30]. Although a previous study reported that MK3 expression was not increased in MK2-deficient cells [31], residual MK3 activity could have effects in *Mapkapk2*^-/-^ mice. Indeed, phenotypic differences between *Mapkapk2*^-/-^ and *Mapkapk2^-/-^Mapkapk3*^-/-^ mice have been documented [32].

A growing body of evidence, including causation studies in mice and correlative studies in human disease tissues, supports a role for MK2 activation in the pathogenesis of chronic inflammatory diseases, including CVD [20]. In addition to the current study and the aforementioned atherosclerosis study using *Mapkapk2*^-/-^ mice [14], other studies have shown that MK2 inhibition can protect against ischemia/reperfusion injury and cardiac hypertrophy, [33, 34], including in a diabetic setting [10]. Most of these studies have used either genetic targeting of MK2 or ATP-competitive MK2 inhibitors, which often cross-react with other kinases, and the cardiac benefits of MK2 inhibition in these studies were not linked to benefits in metabolism. We have shown that a previously developed non-ATP-competitive allosteric MK2 inhibitor with high specificity, cmpd 28, can improve glucose metabolism in obese mice [9], and now we show that two new inhibitors have benefits for both metabolism and atherosclerosis. This demonstration that allosteric MK2 inhibitors can have combined metabolic and vascular benefits in preclinical models should help advance their testing in cardiometabolic disease in humans.

## ACKNOWLEDGMENTS

This work was supported by NIH grants HL087123 (to I.T. and L.O.), DK124457 (to L.O), K99 HL145131 (to A.Y), and an American Heart Association postdoctoral fellowship (20POST35210962) to CK. L.O., B.H., M.H.S.-W., and I.T. are members of Tabomedex Biosciences, Inc., which is developing allosteric MK2 inhibitors for the treatment of T2D.

---

**Figure S1.**
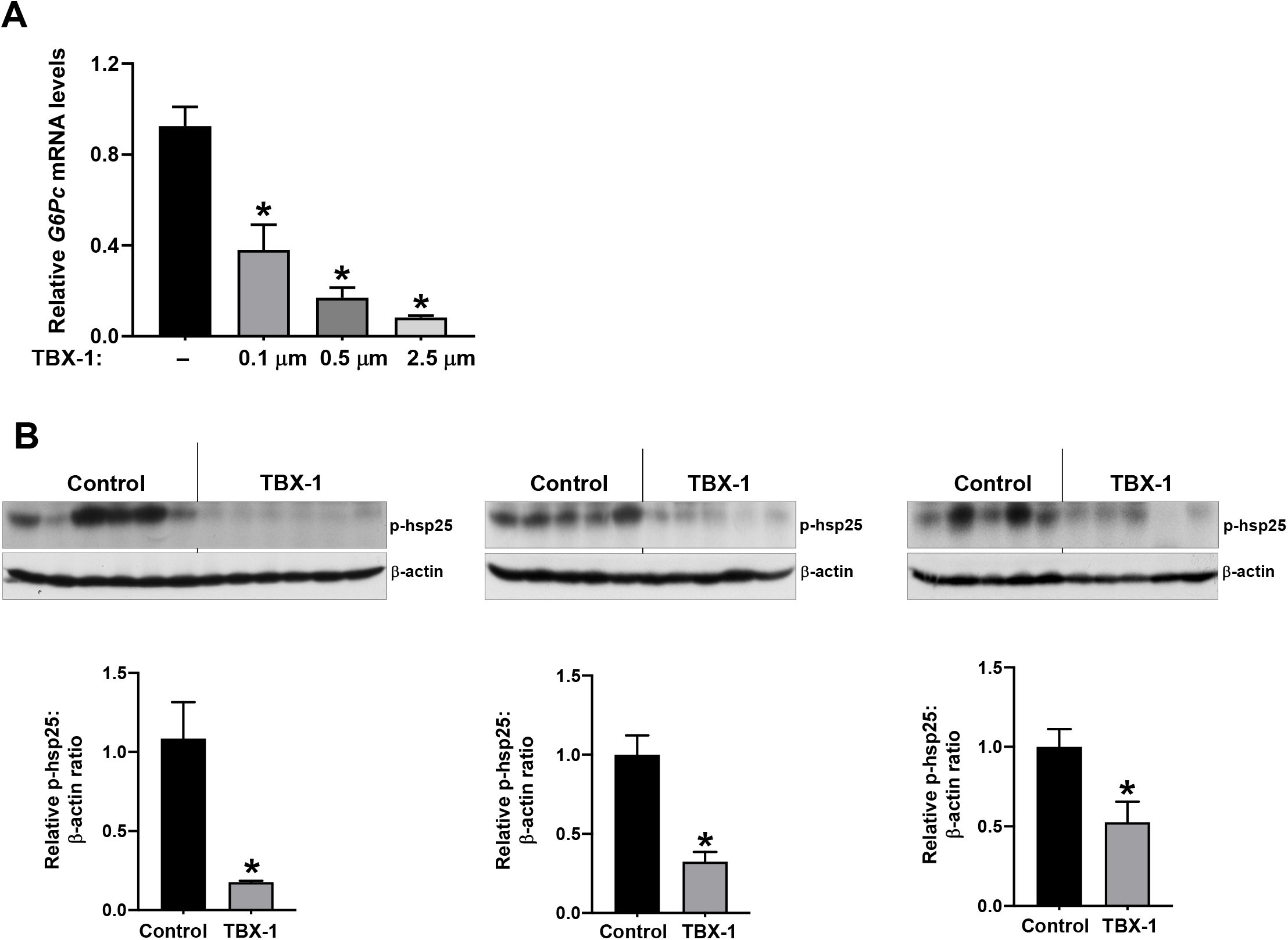
MK2 inhibitor reduces *G6Pc* mRNA in isolated hepatocytes and lowers p-hsp25 levels in WD-fed *Ldlr*^-/-^ mice liver. (**A**) Primary mouse hepatocytes were treated with the indicated concentrations of TBX-1 for 1 h, followed by treatment with TBX-1 and forskolin for 4 h in serum-free media. RNA was assayed for *G6pc* mRNA by RT-qPCR (n = 3; mean ± SEM, p < 0.05). (**B**) Liver extracts from three different randomly selected subsets of TBX-1–treated *Ldlr*^-/-^ mice were assayed for phopsho-hsp25 and β-actin by immunoblot. Densitometric quantification of the data are shown in the bar graphs (n = 5-6 mice/group; mean ± SEM, p < 0.05).

**Figure S2.**
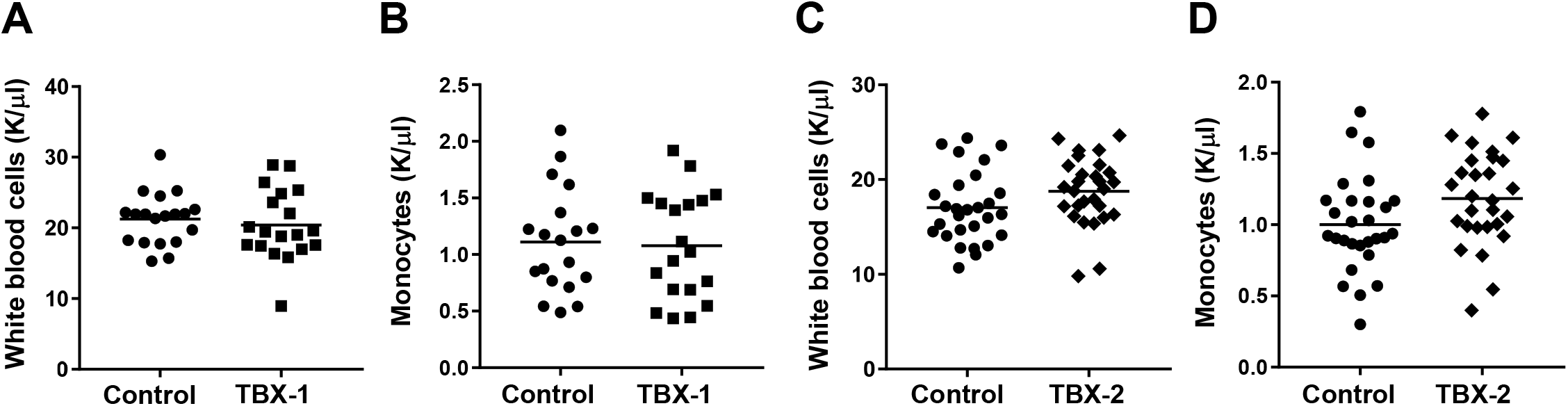
Blood leukocyte and monocyte counts are similar between control and MK2 inhibitor–treated mice. Mice were treated as in Figure 1. Whole blood was analyzed for the number of total white blood cells (**A**) and monocytes (**B**) from a randomly selected subset of TBX-1–treated *Ldlr*^-/-^ mice (n = 19 mice/group; mean ± SEM, p < 0.05). (**C-D**) Same as in (A-B) except that the blood of TBX-2–treated mice was assayed (n = 28 mice/group; mean ± SEM, p < 0.05).

